# Forecasting the numbers of disease vectors with deep learning

**DOI:** 10.1101/2022.11.22.517519

**Authors:** Ana Ceia-Hasse, Carla A. Sousa, Bruna R. Gouveia, César Capinha

## Abstract

Arboviral diseases such as dengue, Zika, chikungunya or yellow fever are a worldwide concern. The abundance of vector species plays a key role in the emergence of outbreaks of these diseases, so forecasting these numbers is fundamental in preventive risk assessment. Here we describe and demonstrate a novel approach that uses state-of-the-art deep learning algorithms to forecast disease vector numbers. Unlike classical statistical and machine learning methods, deep learning models use time series data directly as predictors and identify the features that are most relevant from a predictive perspective. We demonstrate the application of this approach to predict temporal trends in the number of *Aedes aegypti* mosquito eggs across Madeira Island for the period 2013 to 2019. Specifically, we apply the deep learning models to predict whether, in the following week, the number of *Ae. aegypti* eggs will remain unchanged, or whether it will increase or decrease, considering different percentages of change. We obtained high predictive accuracy for all years considered (mean AUC = 0.92 ± 0.05 sd). We also found that the preceding numbers of eggs is a highly informative predictor of future numbers. Linking our approach to disease transmission or importation models will contribute to operational, early warning systems of arboviral disease risk.

## 1. Introduction

Arboviral diseases such as dengue, Zika, chikungunya or yellow fever are a worldwide concern, with large health, social and economic costs. More than half of the world’s population is at risk of disease and economic costs are large, with the costs of dengue alone estimated at US$ 9 billion/year (Mayer et al., 2017; Messina et al., 2019; Shepard et al., 2016). Such diseases are transmitted by mosquitoes of the genus *Aedes* whose distribution is expanding - and projected to expand further in the future - due to climate change and increased transportation and human mobility (Kraemer et al., 2019; Salami et al., 2020b; Santos et al., 2022). These species are now established in many regions of the world, including in several areas of Europe (Capinha et al., 2014; Oliveira et al., 2021), where outbreaks of *Aedes*-borne diseases occurred recently (Brady and Hay, 2019; Sousa et al., 2012). There are no widespread or effective vaccines or treatment for most of these diseases yet, and the spread of *Aedes* spp. calls for integrated tools to predict their presence and number in space and time, to inform timely control initiatives.

Accurate forecasts of *Aedes* spp. numbers are critical to inform public health decision making and implement preventive measures. Since the abundance of vector species plays a key role in disease outbreak emergence (e.g., Jupille et al., 2016; Li et al., 2019; Messina et al., 2019), forecasting such numbers is essential for preventive risk assessment. Most forecasts of mosquito numbers are performed with “classical” statistical and machine learning models, such as generalized linear models *sensu lato* (e.g., generalized additive models, generalized linear mixed models). These models use tabular-type data (i.e., a one-to-one relationship between values of dependent and predictor variables), which implies that the temporal variation in the predictors must be discretized prior to the modelling. This is often performed by applying generic statistical functions (e.g., means or sums) across fixed temporal windows (e.g., weeks, months, years) (e.g., Cheng et al., 2020; da Cruz Ferreira et al., 2017; Li et al., 2019; Monaghan et al., 2019; Poh et al., 2019; Ripoche et al., 2019). Despite its widespread use, this approach it is likely sub-optimal in a high number of cases as there is no guarantee that the resulting set of predictors will be the most adequate from a predictive perspective. This sub-optimality concerns both the statistical functions used to summarize the data and the time scales represented. Recent studies have identified how even simple changes in these parameters can have a strong impact in the quality of the predictions (e.g., Poh et al., 2019).

Here we describe and demonstrate the application of a deep learning-based approach, as an alternative to the classical approaches (e.g., Cheng et al., 2020; da Cruz Ferreira et al., 2017; Li et al., 2019; Monaghan et al., 2019; Poh et al., 2019; Ripoche et al., 2019) for predicting disease vector numbers. We present a novel approach to forecast disease vector numbers that uses state-of-the-art deep learning algorithms and tools (Capinha et al., 2021; Van Kuppevelt et al., 2020). We apply it to predict temporal trends in *Aedes aegypti* mosquito egg numbers. Our major innovation, compared with the state of the art in the field, is that most forecasts of mosquito numbers are performed with “classical” statistical and machine learning models (e.g., Cheng et al., 2020; da Cruz Ferreira et al., 2017; Li et al., 2019; Monaghan et al., 2019; Poh et al., 2019; Ripoche et al., 2019). To our knowledge, this is the first time that time series classification with deep learning is used for modelling the variation of disease vector numbers.

Deep neural networks are artificial neural networks (ANN; Lek and Guégan, 1999; Olden et al., 2008) composed of a large number of trainable parameters (LeCun et al., 2015), which are able to perform complex tasks with high performance, such as computer vision and natural language processing but also image and sound data classification (Christin et al., 2019; LeCun et al., 2015). In the context of disease vector abundance prediction, a key difference of deep learning models over conventional approaches is that the former allow using time series directly as predictors and the relevance of the features found in these data is evaluated by the models themselves (Capinha et al., 2021; Fawaz et al., 2019). Importantly, this relevance (both in terms of the type of data transformation to perform and the temporal extent considered) is guided by the capacity of the models to accurately predict the dependent variable. In other words, deep learning models specifically aim at identifying the set of time series features that most accurately predict disease vector numbers, as opposed to conventional models, which rely on human expertise for this purpose.

As case study, we use the yellow fever mosquito (*Aedes aegypti*) in the Madeira Island, Portugal. This species was first found on this island in 2005 (Margarita et al., 2006) and from late 2012 to early 2013 it was responsible for an outbreak of dengue fever, the first ever in Madeira and the first recorded in Europe since 1927, which infected more than 2000 people (Lourenço and Recker, 2014; Sousa et al., 2012). Due to strong socio-economic relations with South America (namely Brazil and Venezuela), the importation of arbovirus from tropical regions has been considered highly probable (Lourenço and Recker, 2014; Seixas et al., 2019), making the occurrence of future outbreaks of this and other arboviruses possible (Lourenço and Recker, 2014). Because of this, the numbers of *Ae. aegypti* across the island have been closely monitored by local health authorities in recent years and are a key indicator of disease outbreak risk.

We describe and analyse a deep learning-based approach that provides accurate week-ahead forecasts of change in the numbers of *Ae. aegypti* in the Madeira Island. We use time series of the species abundance and of weather variables as predictors and integrate these into a standardized modelling workflow, which could be adapted to provide operational, real-time forecasts, and thus applied in early warning systems of disease risk.

## 2. Material and methods

### 2.1. Mosquito abundance data

Mosquito abundance data were made available by the Regional Health Direction of the Autonomous Region of Madeira (*Direção Regional da Saúde;* https://www.madeira.gov.pt/drs/), the regional health authority responsible for the entomological and epidemiological surveillance in the archipelago. These data originated from 140 ovitraps placed in different locations across the Madeira Island, including air- and seaports, schools, health units and other public and private places, from 2013 to 2019 (Fig 1). This network of traps is monitored weekly, recording the number of eggs of *Ae. aegypti* in each trap.

**Fig 1.**
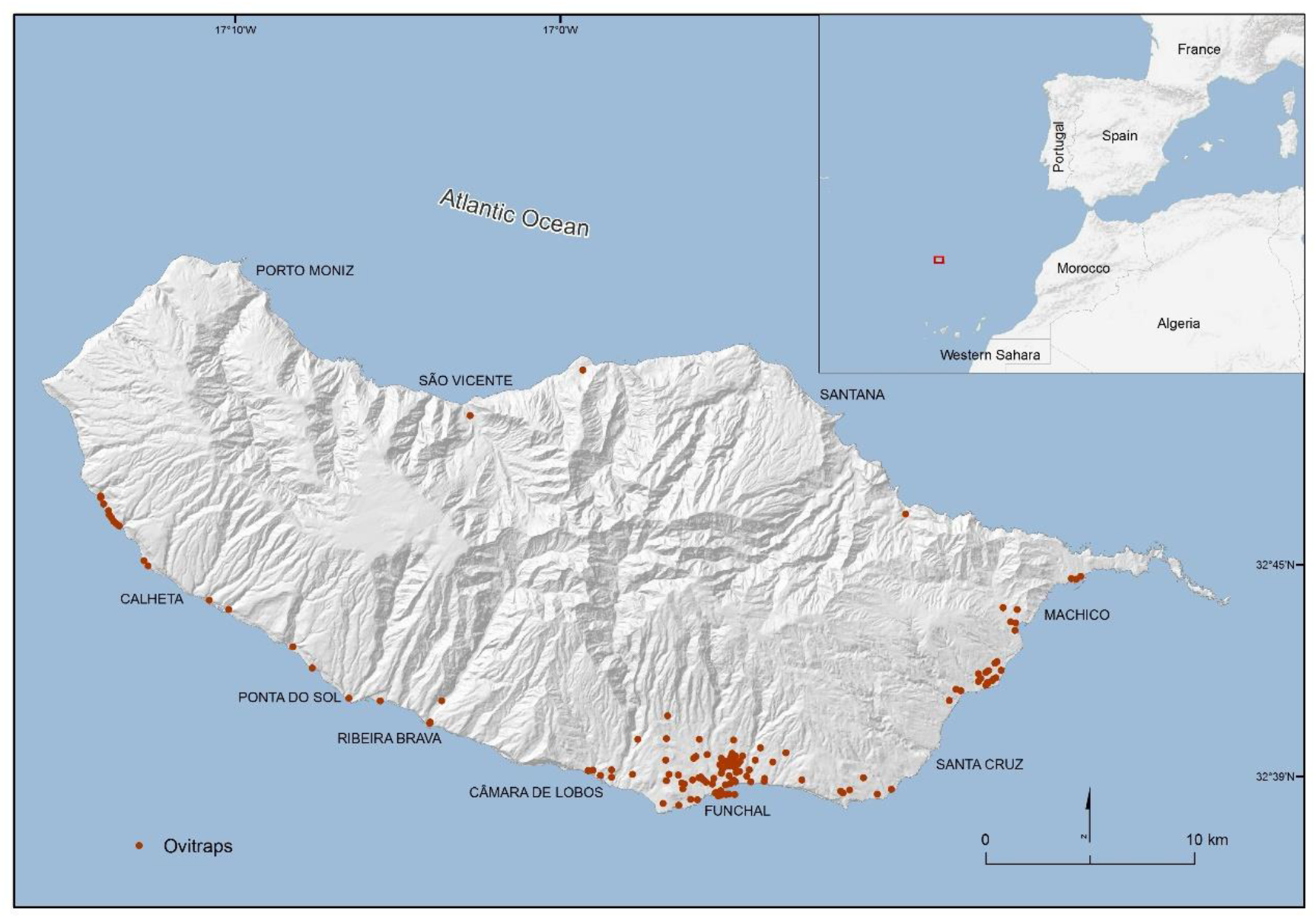
Location of the ovitraps in the Madeira island.

**Fig 2.**
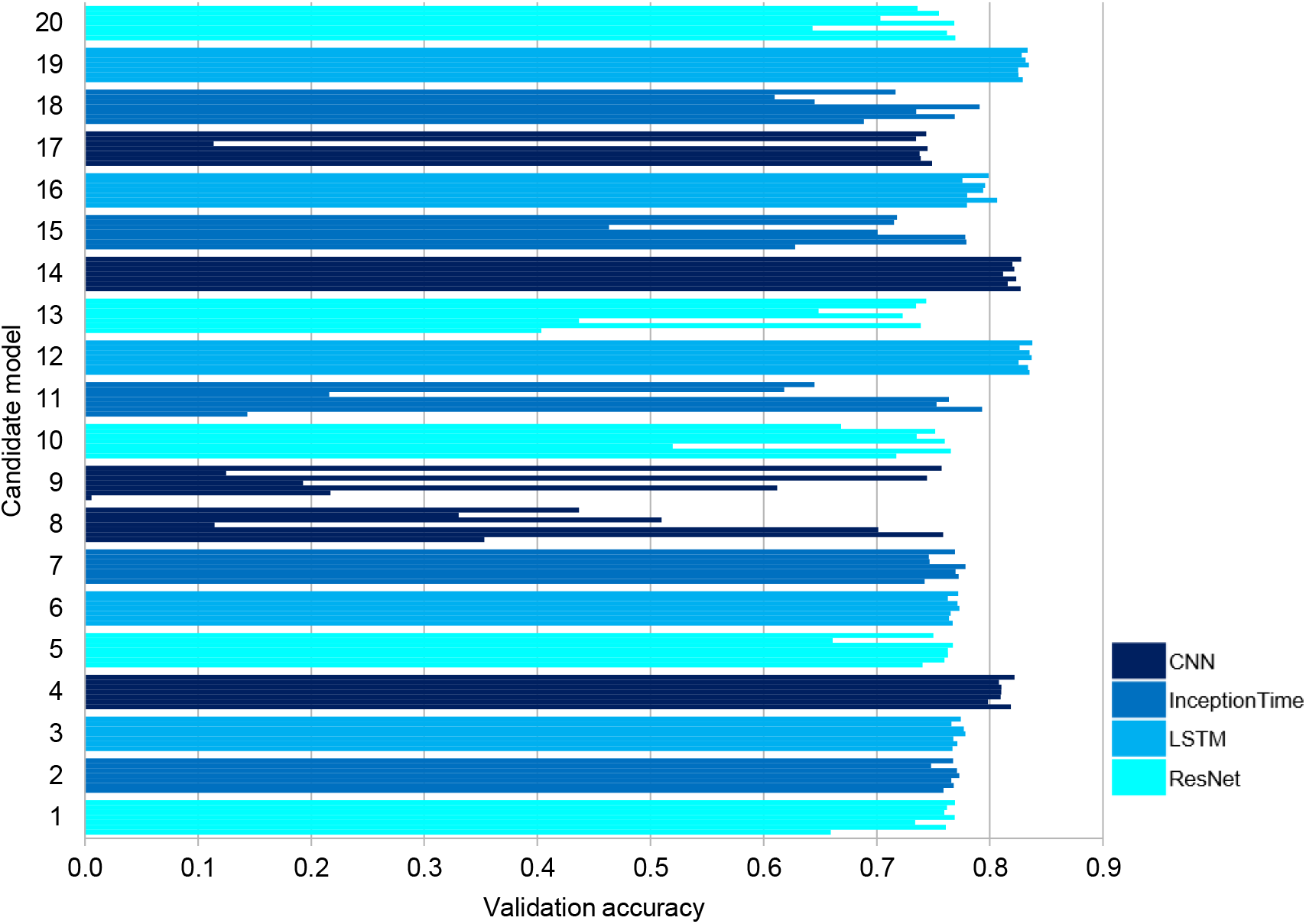
Validation accuracy for candidate models. For each candidate model (1-20 on the vertical axis), each bar corresponds to one test year (from top to bottom: 2019 to 2013). The best model in all years is model 12, having a DeepConvLSTM architecture.

### 2.2. Environmental predictors

As environmental predictors of change in the number of eggs of *Ae. aegypti,* we considered the following weather variables at a daily resolution: mean (*Tmean*), maximum (*Tmax*) and minimum (*Tmin*) temperature, mean (*RHmean*), maximum (*RHmax*) and minimum (*RHmin*) relative humidity, and accumulated precipitation (*Pre).* These variables were selected because of their known role in driving population dynamics of mosquito species (e.g., Cheng et al., 2020; da Cruz Ferreira et al., 2017; Li et al., 2019; Monaghan et al., 2019; Ripoche et al., 2019). Data for these variables was obtained from local weather stations and made available by the Portuguese Institute for Sea and Atmosphere (*Instituto Português do Mar e da Atmosfera;* www.ipma.pt).

### 2.3. Forecasting changes in the number of eggs of Aedes aegypti

We aimed at forecasting relative changes in the numbers of *Ae. aegypti* eggs instead of their absolute numbers. This a common strategy in risk assessment frameworks (e.g., Carbajo and Vezzani, 2015; Tsuda et al., 2016; Vanwambeke et al., 2011). Accordingly, for each trap and observation week, we classified the variation in the number of eggs into one of 7 classes: no change in the number of eggs; decrease between 1 and 25%; decrease between >25 and 50%; decrease >50%; increase between 1 and 25%; increase between >25 and 50%, and increase >50%.

In addition to the weather predictors (see above), we also used the preceding values of egg numbers as predictor of the response variable. To match with the daily resolution of the weather predictors, we replicated the weekly number of eggs 7 times (i.e., one per each day of the corresponding week).

All predictors represented the six-month period that preceded the week of the observation. In other words, the classes of change in number of eggs were modelled as a function of variation in the predictor variables up to 6-months into the past.

This kind of modelling exercise, where time series data is used to predict one of two or more classes, falls withing the scope of time series classification (Capinha et al., 2021; Keogh and Kasetty, 2003). Several deep learning architectures can be used for time series classification, differing in the type of layers they have and on how information flows between them (Fawaz et al., 2019). Because we have no *a priori* reason to expect that a specific deep learning architecture will perform better than others, we evaluated a large number of models having distinct architectures and parametrizations (the so called ‘automated machine learning’ or ‘autoML’ approach; Guyon et al., 2019; Van Kuppevelt et al., 2020).

For this purpose, we use *mcfly* (Van Kuppevelt et al., 2020), a *python* library that allows testing four deep learning architectures frequently used for time series classification: Convolutional Neural Networks (CNN), which have been mostly used for image pattern recognition (Brodrick et al., 2019; Wäldchen and Mäder, 2018) but that are also suited for time series classification (Zhao et al., 2017); Deep Convolutional Long Short-Term Memory networks (DeepConvLSTM), combining convolutional with LSTM recurrent neural networks (Chung et al., 2014), which were developed for sequence-type input data such as time series (Fawaz et al., 2019); Residual Networks (ResNet), which were proposed for image recognition (He et al., 2016) but used recently for time series classification with good performance (Fawaz et al., 2019); and Inception Time networks (InceptionTime), which are a very recent type of architecture proposed explicitly for time series classification (Fawaz et al., 2019). For more details on these modelling architectures see Capinha et al. (2021) and Van Kuppevelt et al. (2020).

### 2.4. Data partition

The ‘autoML’ workflow in *mcfly* requires using multiple partitions from the full data set of response and predictor variables. Here we followed the partition scheme of Capinha et al. (2021), consisting of four partitions that are used at the various stages of model selection and training, and a fifth partition that is used to assess the predictive performance of the fully trained (‘final’) model. To ensure independence between the data used for model training and the data used for model testing, we used the data of each year (i.e., 2013 to 2019) separately for model testing (partition ‘*T*’), and the data for the remaining years for model training. The aim of this procedure is to mimic an operational setting where data available for previous years is used to train a model that will be employed in the real-time forecasting of changes in the number of eggs for the coming week.

The data used for model training and selection were randomly partitioned into: data for training candidate models (25% of the data; *At*); data for validating candidate models (50%; *Av*); data for training the selected candidate model (75%; *Bt* = *At* + *Av*); validation data to determine the optimal number of epochs to train the selected candidate model (25%; *Bv*). Data partition was performed in R with package *dismo* (Hijmans et al., 2017; R Core Team, 2020).

### 2.5. Model selection procedure

The model selection procedure was performed as follows (Capinha et al., 2021; Van Kuppevelt et al., 2020): we randomly generated 5 models for each of the four available deep-ANN architecture types (20 models in total) and trained each one with a small subset of the training data (data partition *At*) for 4 epochs (an “epoch” corresponds to the complete training dataset being passed forward and backward across the network one time; Capinha et al., 2021). The accuracy of candidate models, as provided by *mcfly* (i.e., the “proportion of cases correctly classified”), was then compared using a left-out validation data set (data partition *Av*) and the model with the highest performance was selected for training on the full training data (data partition *Bt*; *Bt* = *At* + *Av*) for up to 30 epochs.

We identified the optimal number of training epochs using data partition B*v*. A too low number of epochs may result in underfitting of the model and thus in reduced predictive performance (since important patterns in the data may be missed), while an excessive number of epochs may result in overfitting of the model, which will reduce its generality and ability to classify new data. The predictive performance of the model at each training epoch each was assessed by measuring the area under the receiving operating characteristic curve (AUC), which is usually used in ecology and is not influenced by differences in the prevalence of classes (Dyderski et al., 2018). AUC was computed within R, using package *cvAUC* (LeDell et al., 2014; R Core Team, 2020).

Finally, the performance of the model trained with the optimal number of epochs was evaluated using a test data set (data set *T*), using AUC as the accuracy metric as well. This procedure was repeated so that each year was used as test data (as partition ‘*T*’).

### 2.6. Importance of predictors

We assessed the importance of each predictor variable by performing simulations with randomized test data, where in each simulation one predictor variable was randomized at a time, while keeping the others unchanged (Molnar, 2020), and comparing the AUC from such simulations with the AUC from the original models. We did this for each different test year. We prepared the randomized data in R (using function *sample* from package *base*) (R Core Team, 2020).

See Appendix A for example data and code for implementing data importation and modelling procedures. For more details on the deep learning techniques we used please see Capinha et al. (2021) and Van Kuppevelt et al. (2020).

## 3. Results

Regarding the percentage of observations per class of variation in the number of eggs between subsequent weeks, the class with more observations was “no change”, in all the years (table 1); these correspond mainly to cases where the number of eggs was continuously zero. The second and third most represented classes correspond to the largest (>50%) decrease and increase (respectively), and the classes the least represented correspond to the lower decreases or increases in the number of eggs (i.e., 1 to 25% and >25 to 50%; table 1).

**Table 1.** Percentage of observations per class of change in number of eggs between subsequent weeks in each year. “inc.” stands for increase and “dec.” stands for decrease in the number of eggs.

|  | 2013 | 2014 | 2015 | 2016 | 2017 | 2018 | 2019 |
| --- | --- | --- | --- | --- | --- | --- | --- |
| no change | 81.7 | 78.0 | 77.6 | 74.7 | 74.9 | 79.4 | 75.3 |
| inc. 1-25% | 0.7 | 0.6 | 0.7 | 1.0 | 0.6 | 0.7 | 0.8 |
| inc. 25-50% | 0.8 | 1.0 | 1.4 | 1.2 | 1.6 | 1.0 | 0.9 |
| inc. >50% | 7.4 | 8.9 | 9.1 | 10.4 | 10.3 | 8.4 | 10.4 |
| dec. 1-25% | 0.6 | 0.6 | 0.8 | 0.9 | 0.8 | 0.5 | 0.7 |
| dec. 25-50% | 0.3 | 0.7 | 0.7 | 0.4 | 0.5 | 0.3 | 0.5 |
| dec. >50% | 8.5 | 10.1 | 9.8 | 11.5 | 11.4 | 9.6 | 11.4 |

Concerning the performance of the candidate models, the best validation accuracies were achieved by model 12 in all the years (accuracy = 0.833 ± 0.005; mean of years ± sd) (Fig 2), a model having a Deep Convolutional Long Short-Term Memory (DeepConvLSTM) architecture.

Model 19, also with a DeepConvLSTM architecture had high validation accuracies too (accuracy = 0.829 ± 0.003; mean of years ± sd), followed by model 14, with a Convolutional Neural Network (CNN) architecture and validation accuracy of 0.821± 0.005 (mean of years ± sd).

After full training and identification of the optimal number of training epochs, this model delivered an excellent average predictive performance for all years (mean AUC = 0.92 ± 0.05 sd). Only in a few cases the performance went below this threshold, namely for class “increase >50%” where performance was fair (i.e., AUC from 0.7 to 0.8) for years 2013 and 2016, and good (AUC 0.8 to 0.9) for the remaining years, and for class “increase 25-50%” for year 2017, where the model performance was good (AUC = 0.88) (table 2).

**Table 2.** AUC obtained in simulations with the original models: mean (and standard deviation) of classes and AUC per class. The year used for testing model performance is given in the column headers. “inc.” stands for increase, “dec.” stands for decrease.

|  | 2013 | 2014 | 2015 | 2016 | 2017 | 2018 | 2019 |
| --- | --- | --- | --- | --- | --- | --- | --- |
| **mean** | 0.92 | 0.93 | 0.92 | 0.90 | 0.91 | 0.91 | 0.92 |
| **(sd)** | (0.06) | (0.04) | (0.04) | (0.06) | (0.05) | (0.05) | (0.05) |
| **no change** | 0.92 | 0.94 | 0.94 | 0.91 | 0.93 | 0.92 | 0.92 |
| **inc. 1-25%** | 0.95 | 0.94 | 0.93 | 0.91 | 0.92 | 0.92 | 0.94 |
| **inc. 25-50%** | 0.93 | 0.93 | 0.93 | 0.90 | 0.88 | 0.92 | 0.94 |
| **inc. >50%** | 0.78 | 0.83 | 0.83 | 0.78 | 0.81 | 0.80 | 0.80 |
| **dec. 1-25%** | 0.95 | 0.96 | 0.94 | 0.94 | 0.94 | 0.93 | 0.95 |
| **dec. 25-50%** | 0.96 | 0.94 | 0.94 | 0.93 | 0.95 | 0.96 | 0.93 |
| **dec. >50%** | 0.94 | 0.95 | 0.96 | 0.94 | 0.93 | 0.94 | 0.93 |

Concerning the importance of variables, AUC decreased most notoriously when the predictor ‘number of eggs’ was randomized, a result observed for all test years (Fig 3). Slight decreases in performance are also apparent when temperature was randomized (either mean, maximum or minimum) for most test years.

**Fig 3.**
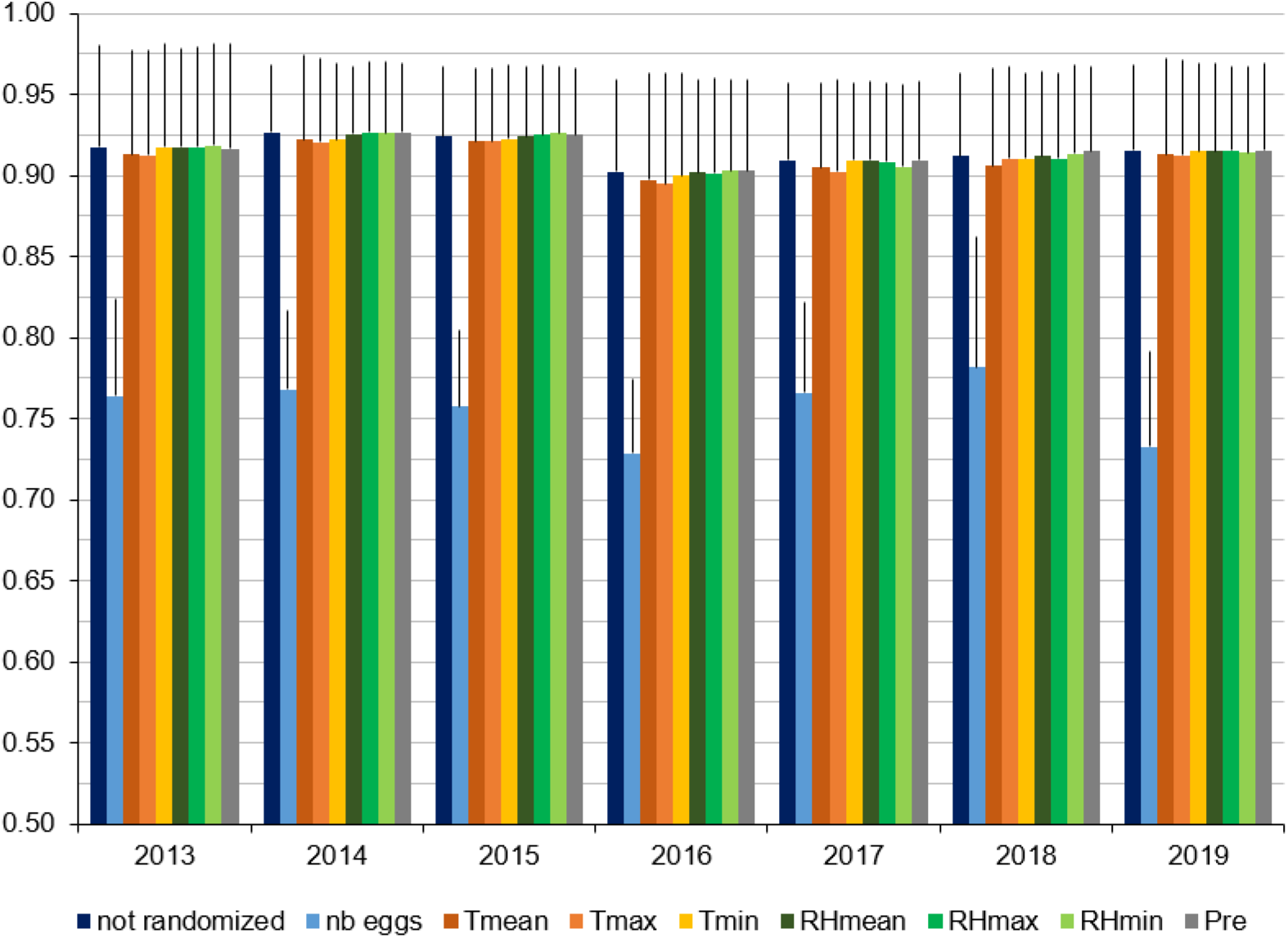
AUC (mean of classes and standard deviation) for simulations performed with original (not randomized) and randomized predictor variables. (e.g., the columns “nb eggs” correspond to the simulations performed with randomized number of eggs, the columns “*Tmean*” correspond to the simulations performed with randomized mean temperature, and so on), for each year used as test data (indicated in the horizontal axis). *Tmean*, *Tmax*, *Tmin*: mean, maximum and minimum temperature (respectively), *RHmean*, *RHmax*, *RHmin*: mean, maximum and minimum relative humidity (respectively), *Pre*: precipitation.

## 4. Discussion

Obtaining accurate predictions of disease vector distribution and numbers is critical to inform decision making and implement timely and effective control actions, in line with the United Nations Sustainable Development Goals 3 and 13 of ending vector-borne disease epidemics and strengthening early warning capacity of global health risks (UN General Assembly, 2015).

In this work we used a deep learning approach to forecast the numbers of *Ae. aegypti* in Madeira island. To our knowledge, this is the first time that time series classification with deep learning is used for predicting variation of disease vector numbers. We also showed that the predictive accuracy of the approach was high, with the cross-class performances being good to excellent for all years considered.

We provide the code and data files we used as appendix for those who which to replicate our study. Validation was conducted by assessing the performance of the models using seven different data sets and time periods independently, providing independent validation and demonstrating the advantages of the approach, given the high predictive accuracy obtained.

We selected the candidate model with the highest validation accuracy in all years as the final model to train with the full training data set and obtain the predictions. However, other models also achieved high validation accuracies. Future work could include testing whether making predictions with an ensemble of such models would increase the robustness of predictions. However, for the purpose of demonstrating our approach we considered appropriate to use only the model with the highest validation accuracy.

Our results are supportive of a wider testing and application of deep learning for predicting disease vector numbers. A defining feature of this approach is that it allows using raw time series data as predictors, being able to automatically identify relevant information such as thresholds and lag effects (Fawaz et al., 2019; Ryo et al., 2019) that could otherwise be missed when using classical approaches, which rely on temporally aggregated (and thus simplified) variables (Capinha et al., 2021). This capacity of deep learning models can be an important asset in face of the growing availability of time series data, namely from mosquito monitoring programs and environmental sensors (e.g., satellite imagery and meteorological stations; Reichstein et al., 2019), that can be directly fed into the models (Capinha et al., 2021).

We also identified that the number of eggs in the previous weeks was the most important predictor of variation in number of eggs, in agreement with previous studies (da Cruz Ferreira et al., 2017), and which suggests this variable has a high intrinsic predictability (Pennekamp et al., 2019). The relative importance of individual predictors representing other factors (temperature, relative humidity, and precipitation) was lower, despite an apparent relevance being also detected for temperature-related time series. We note that in the case of temperature and relative humidity, this apparently lower relevance may also result from correlations between the three measurements used to represent these factors (i.e., minimum, mean and maximum values). In other words, the models may be obtaining relevant information from time series representing a different measurement of the same factor.

The approach presented here did not consider spatial structure explicitly. Given that the previous number of eggs had high explanatory power, the past number of eggs in nearby traps as predictors could also be included. We could also consider a leave one trap out method of validation, to see if the developed models are general spatially.

Our models use tabular-type data and thus the temporal variation in the predictors was discretized prior to modelling. Although not the objective of our work, there are statistical models that can estimate lags between time series and disease outcomes, allowing the exploration of temporal lags without specifying lags a priori (e.g., Davis et al., 2018; Smith et al., 2020). Additionaly, longer time series could also be considered as this has been reported for infectious disease predictions (e.g., Smith et al., 2020).

Deep learning is a promising avenue but there is room for improvement and limitations to be mentioned and addressed in future studies. On one hand, the approach presented can be adapted for other specific classification tasks, and model generation and selection in *mcfly* can be fine-tuned. For example, the selection of candidate models can be adjusted to generate only one type of model architecture (Capinha et al., 2021; Van Kuppevelt et al., 2020).

Also, deep learning approaches could be compared with the more classical statistical and machine learning approaches to assess how model performances differ between methods. Our study design could be complemented with a comparison with a simple baseline model and with “classical” statistical and machine learning models, such as generalized linear models *sensu lato* (e.g., Li et al., 2019; Poh et al., 2019; Ripoche et al., 2019). However, our objective was to show our approach works, which is clearly demonstrated by the high predictive accuracy obtained. Furthermore, the ecological interpretability of deep learning models and their outputs should be improved (Reichstein et al., 2019). This is being now tackled by the field of “explainable artificial intelligence” to understand how the models work and explain their outcomes (Shickel and Rashidi, 2020; Siddiqui et al., 2019).

Despite these challenges, we presented a well performing modelling framework that can be used by non-experts in deep learning (Capinha et al., 2021; Van Kuppevelt et al., 2020) to predict disease vector numbers. Our work has a clear public health relevance. Integrating this approach into existing disease transmission or importation models (Lieberthal and Gardner, 2021; Salami et al., 2020b, 2020a) or with disease-related Internet search activity (Aiken et al., 2020; Yang et al., 2017) and using high resolution environmental data will likely contribute to improve operational, early warning systems of disease risk and guide the implementation of mosquito control measures.

## Supporting information

CeiaHasseetal_Supplementary_material

## Acknowledgements

We thank the personnel of the Regional Health Direction of the Autonomous Region of Madeira (*Direção Regional da Saúde*) involved in egg collection and analysis and the Portuguese Institute for Sea and Atmosphere (*Instituto Português do Mar e da Atmosfera*), namely Dr. Victor Prior, for the weather data. Maurício Santos drew Fig 1. ACH, CAS and CC were supported by Portuguese National Funds through Fundação para a Ciência e a Tecnologia (ACH and CAS: PTDC/SAU-PUB/30089/2017 and GHTM-UID/Multi/04413/2020; CC: CEECIND/02037/2017, UIDB/00295/2020 and UIDP/00295/2020). The funders had no role in study design, data collection and analysis, decision to publish, or preparation of the manuscript.

## Appendix A. Supplementary material

**S1 Text. Code for implementing data importation and modelling procedures.**

**S1 Data. At classes eggs.** Data file with egg classes from *At* partition.

**S1 Data. At nb eggs.** Data file with number of eggs from *At* partition.

**S1 Data. At prec.** Data file with precipitation from *At* partition.

**S1 Data. At rhmax.** Data file with maximum relative humidity from *At* partition.

**S1 Data. At rhmean.** Data file with mean relative humidity from *At* partition.

**S1 Data. At rhmin.** Data file with minimum relative humidity from *At* partition.

**S1 Data. At tmax.** Data file with maximum temperature from *At* partition.

**S1 Data. At tmean.** Data file with mean temperature from *At* partition.

**S1 Data. At tmin.** Data file with minimum temperature from *At* partition.

**S2 Data. Av classes eggs.** Data file with egg classes from *Av* partition.

**S2 Data. Av nb eggs.** Data file with number of eggs from *Av* partition.

**S2 Data. Av prec.** Data file with precipitation from *Av* partition.

**S2 Data. Av rhmax.** Data file with maximum relative humidity from *Av* partition.

**S2 Data. Av rhmean.** Data file with mean relative humidity from *Av* partition.

**S2 Data. Av rhmin.** Data file with minimum relative humidity from *Av* partition.

**S2 Data. Av tmax.** Data file with maximum temperature from *Av* partition.

**S2 Data. Av tmean.** Data file with mean temperature from *Av* partition.

**S2 Data. Av tmin.** Data file with minimum temperature from *Av* partition.

**S3 Data. Bt classes eggs.** Data file with egg classes from *Bt* partition.

**S3 Data. Bt nb eggs.** Data file with number of eggs from *Bt* partition.

**S3 Data. Bt prec.** Data file with precipitation from *Bt* partition.

**S3 Data. Bt rhmax.** Data file with maximum relative humidity from *Bt* partition.

**S3 Data. Bt rhmean.** Data file with mean relative humidity from *Bt* partition.

**S3 Data. Bt rhmin.** Data file with minimum relative humidity from *Bt* partition.

**S3 Data. Bt tmax.** Data file with maximum temperature from *Bt* partition.

**S3 Data. Bt tmean.** Data file with mean temperature from *Bt* partition.

**S3 Data. Bt tmin.** Data file with minimum temperature from *Bt* partition.

**S4 Data. Bv classes eggs.** Data file with egg classes from *Bv* partition.

**S4 Data. Bv nb eggs.** Data file with number of eggs from *Bv* partition.

**S4 Data. Bv prec.** Data file with precipitation from *Bv* partition.

**S4 Data. Bv rhmax.** Data file with maximum relative humidity from *Bv* partition.

**S4 Data. Bv rhmean.** Data file with mean relative humidity from *Bv* partition.

**S4 Data. Bv rhmin.** Data file with minimum relative humidity from *Bv* partition.

**S4 Data. Bv tmax.** Data file with maximum temperature from *Bv* partition.

**S4 Data. Bv tmean.** Data file with mean temperature from *Bv* partition.

**S4 Data. Bv tmin.** Data file with minimum temperature from *Bv* partition.

**S5 Data. T classes eggs.** Data file with egg classes from *T* partition.

**S5 Data. T nb eggs.** Data file with number of eggs from *T* partition.

**S5 Data. T prec.** Data file with precipitation from *T* partition.

**S5 Data. T rhmax.** Data file with maximum relative humidity from *T* partition.

**S5 Data. T rhmean.** Data file with mean relative humidity from *T* partition.

**S5 Data. T rhmin.** Data file with minimum relative humidity from *T* partition.

**S5 Data. T tmax.** Data file with maximum temperature from *T* partition.

**S5 Data. T tmean.** Data file with mean temperature from *T* partition.

**S5 Data. T tmin.** Data file with minimum temperature from *T* partition.

**S6 Data. T nb eggs random.** Data file with randomized number of eggs from *T* partition.

**S6 Data. T prec random.** Data file with randomized precipitation from *T* partition.

**S6 Data. T rhmax random.** Data file with randomized maximum relative humidity from *T* partition.

**S6 Data. T rhmean random.** Data file with randomized mean relative humidity from *T* partition.

**S6 Data. T rhmin random.** Data file with randomized minimum relative humidity from *T* partition.

**S6 Data. T tmax random.** Data file with randomized maximum temperature from *T* partition.

**S6 Data. T tmean random.** Data file with randomized mean temperature from *T* partition.

**S6 Data. T tmin random.** Data file with randomized minimum temperature from *T* partition.

